# The basis of a more contagious 501Y.V1 variant of SARS-COV-2

**DOI:** 10.1101/2021.02.02.428884

**Authors:** Haolin Liu, Qianqian Zhang, Pengcheng Wei, Zhongzhou Chen, Katja Aviszus, John Yang, Walter Downing, Shelley Peterson, Chengyu Jiang, Bo Liang, Lyndon Reynoso, Gregory P. Downey, Stephen K. Frankel, John Kappler, Philippa Marrack, Gongyi Zhang

## Abstract

Severe acute respiratory syndrome-related coronavirus 2 (SARS-CoV-2) is causing a world-wide pandemic. A variant of SARS-COV-2 (20I/501Y.V1) recently discovered in the United Kingdom has a single mutation from N501 to Y501 within the receptor binding domain (Y501-RBD), of the Spike protein of the virus. This variant is much more contagious than the original version (N501-RBD). We found that this mutated version of RBD binds to human Angiotensin Converting Enzyme 2 (ACE2) a ~10 times more tightly than the native version (N501-RBD). Modeling analysis showed that the N501Y mutation would allow a potential aromatic ring-ring interaction and an additional hydrogen bond between the RBD and ACE2. However, sera from individuals immunized with the Pfizer-BioNTech vaccine still efficiently block the binding of Y501-RBD to ACE2 though with a slight compromised manner by comparison with their ability to inhibit binding to ACE2 of N501-RBD. This may raise the concern whether therapeutic anti-RBD antibodies used to treat COVID-19 patients are still efficacious. Nevertheless, a therapeutic antibody, Bamlanivimab, still binds to the Y501-RBD as efficiently as its binds to N501-RBD.

---

The SARS-CoV-2 virus has infected over one hundred million people (COVID-19 patients) and caused more than two million deaths to date (WHO_COVID-19_patients 2020). The numbers of affected people continue to grow rapidly, emphasizing the need for rapid use of effective vaccines. Although two mRNA (Pfizer-BioNTech COVID-19 Vaccine and MODERNA respectively) Spike protein based vaccines have been approved for emergency use in the USA (Polack et al. 2020; Baden et al. 2020) the increasing number of Spike variants that have appeared around the world raise concerns about the continued efficacy of the vaccines (SARS-COV-2-Variants 2021). It has been reported that ~90% of broadly neutralizing anti-SARS-CoV-2 antibodies from COVID-19 patients engage the receptor binding domain (RBD) of the virus Spike protein (Piccoli et al. 2020). Monoclonal antibodies specifically targeting the native form of Spike developed by Regeneron and Eli Lilly have been approved by the FDA for emergency use (Baum et al. 2020; Gottlieb et al. 2021; Bamlanivimab 2020; casirivimab and imdevimab 2020). An N501Y variant of SARS-CoV-2 (B.1.1.7, 20I/501Y.V1), first emerging in the United Kingdom and now spreading to the rest of the world recently, appears much more contagious than the original version (SARS-COV-2-Variants 2021). Furthermore, this same N501Y mutation is also found in a variant (B.1.351, 20H/501Y.V2) from South Africa and a variant (P1, 20J/501Y.V3) from Brazil (SARS-COV-2-Variants 2021). Unfortunately, this mutation is located right at the surface of interaction between RBD and ACE2 (Lan et al. 2020). Here we tested whether B.1.1.7, 20I/501Y.V1 might have a higher affinity for ACE2, thus accounting at least in part for the greater infectivity of SARS-CoV-2 expressing this variant. We found that the mutated RBD has a ~10 times fold higher binding affinity for ACE2 than N501-RBD has. Structural modeling data showed that N501-RBD can form an additional aromatic ring-ring interaction and an additional hydrogen bond with ACE2. In spite of this, sera from individuals immunized with the Pfizer-BioNTech vaccine still efficiently block ACE2 binding to Y501-RBD. Structural modelling shows that a new aromatic ring-ring interaction is formed in the content of new interactions with an additional hydrogen bond. However, serums from individuals immunized with Pfizer-BioNTech still efficiently block ACE2 binding to Y501-RBD. Furthermore, Bamlanivimab, the recently FDA approved therapeutic antibody drug to treat COVID-19 patients (Bamlanivimab 2020), still binds to the variant Y501-RBD as efficiently as it binds to N501-RBD RBD giving hope that treatment with the existing monoclonal antibodies may still help COVID patients.

To investigate the basis for the more highly infectious property of the N501Y variant of SARS-CoV-2, we expressed two versions of SARS-CoV-2 RBD, N501-RBD and Y501-RBD (residues 319-541 aa) in 293F cells. Purified proteins of both these types were subjected to Surface Plasmon Resonance binding assays on a Biacore machine to examine their binding affinities for ACE2. The binding affinity between the native form of RBD (N501-RBD) and ACE2 is ~5.76 nM (**Fig. 1A**), an affinity that is similar to that, ~4.5 nM, reported previously (Lan et al. 2020). To our surprise, the binding affinity of the mutated version of the RBD, Y501-RBD increases dramatically, to ~0.566 nM (**Fig. 1B**), an affinity that is almost ~10 times fold higher than that of N501-RBD for ACE2 (**Fig. 1A**). It has previously been reported that the affinity of the RBD of SARS-CoV-2 for ACE2 is ~10 times fold higher than that of the RBD of SARS-CoV for the same ligand, ACE2 (Lan et al. 2020; Wrapp et al. 2020). The ~10 times fold increase of binding affinity could be one of major contributors to the fact that SARS-CoV-2 is more infectious than SARS-CoV (Starr et al. 2020). The current mutation of N501 to Y501 and consequent ~10 times greater affinity for ACE2 may therefore account for the increased rate of infections in the United Kingdom and, likewise, for the increased transmission rate of both the South African (20H/501Y.V2) and Brazil variants (20J/501Y.V3), although their K417N, E484K besides N501Y changes may also contribute (Tegally et al. 2020).

**Figure 1.**
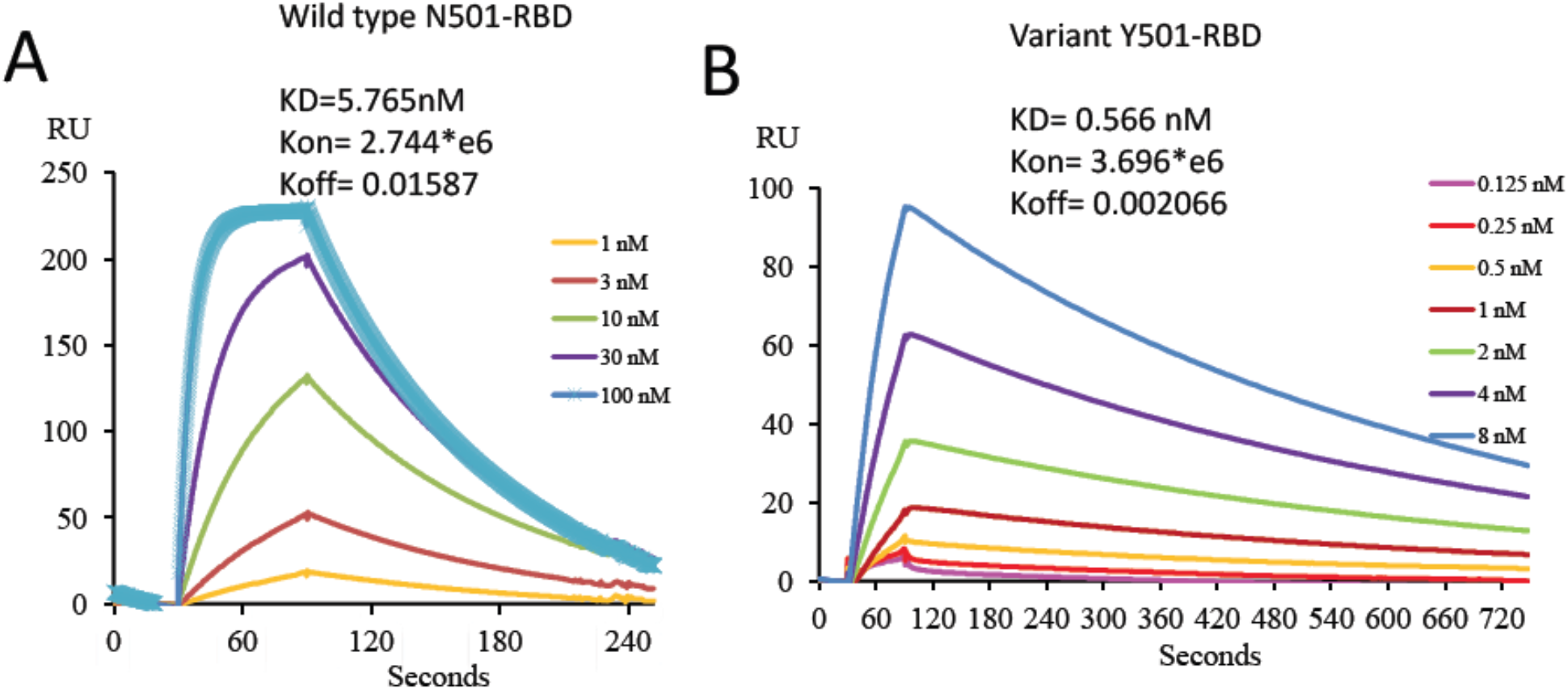
The binding affinities of N501-RBD and mutant Y501-RBD to ACE2 measured on a Biacore machine. **A.** the binding between N501-RBD and ACE2. **B.** Binding between Y501-RBD and ACE2.

A 2.9Å resolution Cryo-EM structure of a predicted potential variant with the highest binding affinity of RBD to ACE2 containing mutations of I358F, V445K, N460K, I468T, T470M, S477N, E484K, Q498R, N501Y with ACE2 has been published on line (Zahradnik et al. 2021), and shows a detailed interaction of the side-chain of Y501 of the RBD and Y41 of ACE2. However, due to the artificial introduction of Q498R (this mutation does not exist in current variants) at this microenvironment around Y501, the structure could not unequivocally attribute the ~10 times fold increase in affinity of Y501-RBD versus N501-RBD for ACE2 to the N501Y mutation. The structure of the RBD bearing just the N501Y mutation and ACE2 has not yet been reported. To compensate for the current absence of the needed structure we used modern molecular modeling programs to make a prediction of the relevant binding sites. Thus, based on the reported structure of the N501-RBD and ACE2 (PDB ID: 6M0J)(Lan et al. 2020), we generated an Y501-RBD and ACE2 complex in program COOT (Emsley and Cowtan 2004a) and used the docking program HADDock 2.2 server to carry out optimization (Van Zundert et al. 2016) (**Fig. 2**). In the case of the wild type SARS-CoV-2 Spike, the side chain of N501, within N501-RBD forms a hydrogen bond with Y41 of ACE2 (**Fig. 2A**). This interaction is of course markedly changed in the interaction between the Y501-RBD mutant and ACE2. Although the hydrogen bond between N501-RBD and Y41 of ACE2 disappears for the Y501-RBD mutant, two new forms of interactions, hydrogen bonds and a hydrophobic interaction are predicted to exist between Y501-RBD and ACE2 (**Fig. 2B**). First, Y501 of Y501-RBD forms two new hydrogen bonds with the side chains of D38 and K353 of ACE2 (**Fig. 2B**). Second, the aromatic ring of Y501 also has a strong aromatic stacking interaction (pi stacking) with the aromatic ring of Y41 of ACE2 (**Fig. 2B**). It is quite likely that the ring-ring interaction may contribute the much higher binding affinity between Y501-RBD and ACE2. However, from the cryo-EM structure, the side chain of R498 interferes with the ring-ring packing, it is possible that two aromatic-rings and side-chain of D38 form a negatively charged cage to hold the side-chain of R498 leading to the ultra-high binding affinity as showed (Zahradnik et al. 2021). Nevertheless, these detailed interactions still need confirmation with a high-resolution structure of Y501-RBD and ACE2.

**Figure 2.**
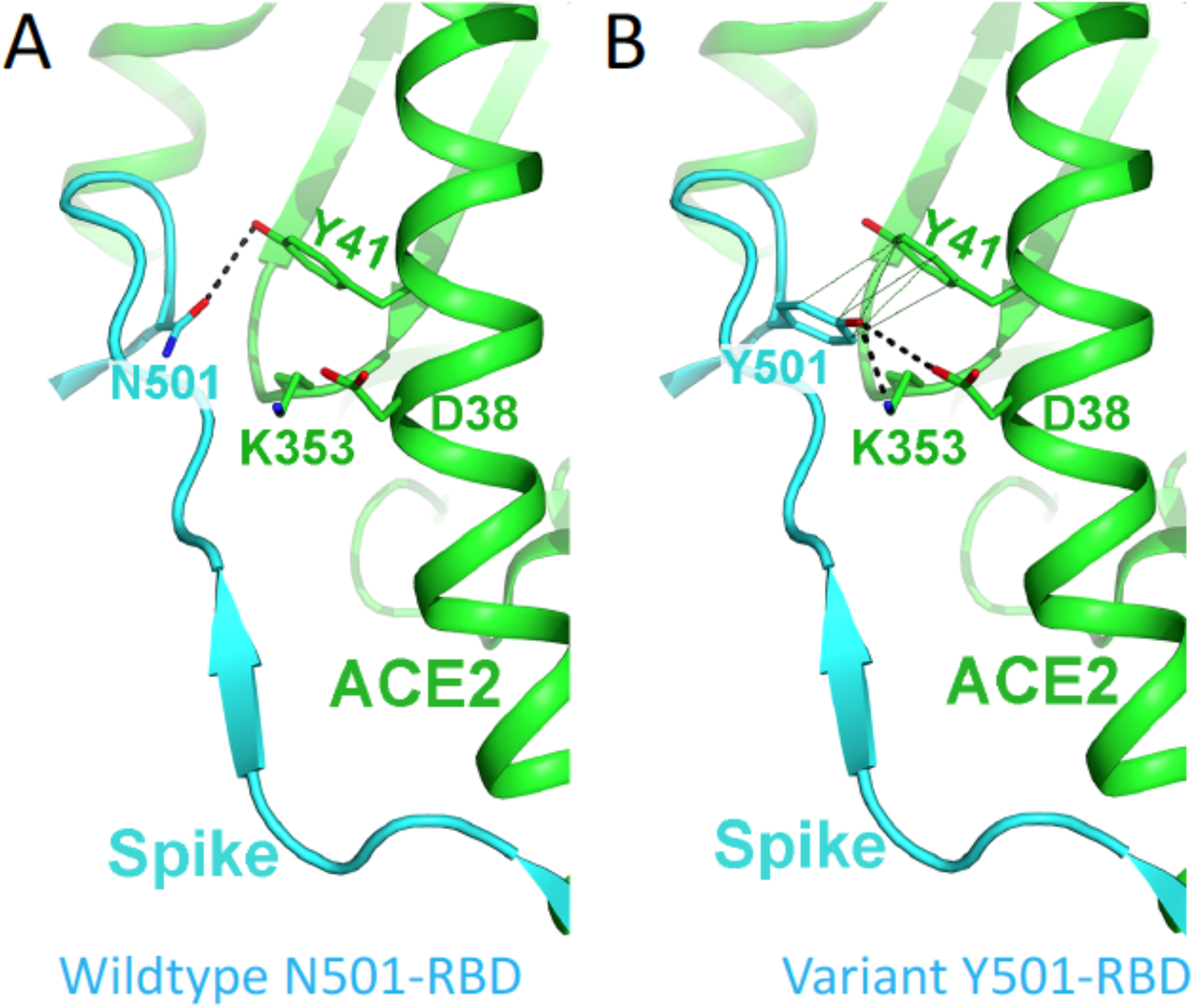
Diagram illustrating the interaction between SARS-CoV-2 RBD and ACE2. **A.** WT, N501-RBD and ACE2. **B.** Y501-RBD and ACE2. Note that the structure of N501Y mutant was mutated from the WT structure (PDB ID: 6M0J). The new hydrogen bonds were shown as heavy dash lines, the aromatic stacking interaction was shown as light dash line.

As it has been reported that ~90% of broad neutralizing antibodies from COVID-19 patients target RBD of the Spike (Piccoli et al. 2020). Thus it is important to test if the binding of these broadly neutralizing antibodies is affected by the change of N501-RBD to Y501-RBD. To investigate this problem we tested sera from individuals immunized with the mRNA vaccine from Pfizer-BioNTech. High titers of antibodies from five immunized individuals were detected by N501-RBD coated ELISA plate (**Fig. S1**), suggesting that these individuals have triggered the expected strong immune response to the RBD after immunization with the vaccine. To test the blocking property of broadly neutralizing antibodies toward both native and mutated RBDs, both RBDs are immobilized on a Biacore Chip and serum from one vaccinated individual was injected at different concentrations to test its ability to block binding of the RBD to ACE2 (**Fig. 3**). The serum blocked binding to ACE2 of both N501-RBD and Y501-RBD (**Fig. 3A and Fig. 3B**). There is no ACE2 binding for RBDs treated with undiluted serum (1* in **Fig. 3A and Fig. 3B)** while barely binding of ACE2 when the serum is diluted 2, 3, 4, and 5 times (2-5* in **Fig. 3A and Fig. 3B)**. These binding data suggest that vaccine from the Pfizer-BioNTech could protect people from both variants of SARS-COV-2. However, the blocking efficacy of the serum for Y501-RBD was slightly compromised with the increasing folds of dilution of the serum (**Fig. 3C**). This slight loss of efficacy could raise concerns for therapeutic antibodies, which are specific for the RBD to block ACE2 binding.

**Figure 3.**
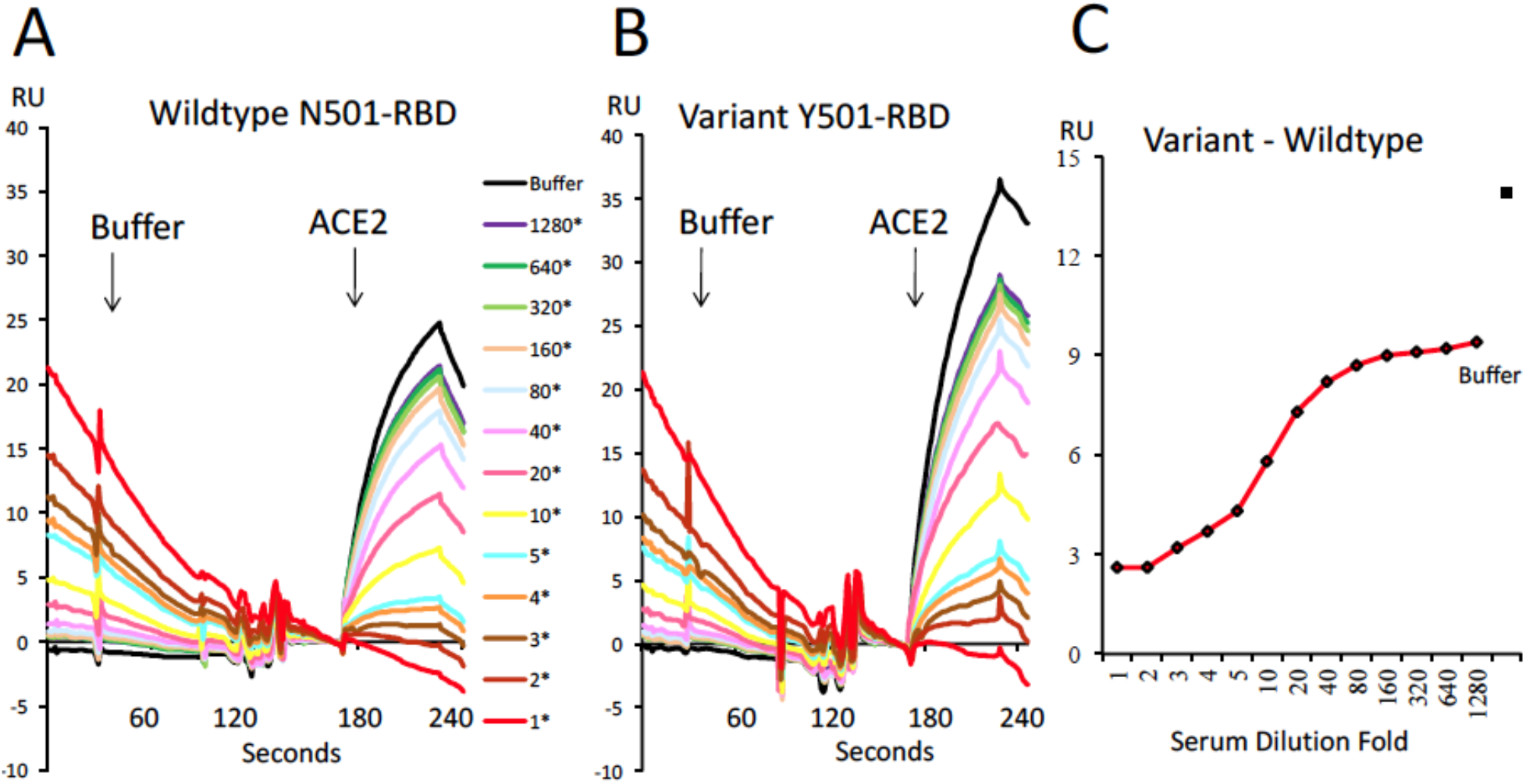
N501-RBD and Y501-RBD treated with vaccinated human serum block ACE2 binding. **A.** The blocking of ACE2 to N501-RBD after treatments with the serum solution with a series of dilutions. Injecting 20 μΙ serum with or without dilution for 1 minutes, washing with buffer (Arrow) with 20 μΙ for 1 minutes, then injecting 20 μΙ of 5 μg/ml ACE2 for 1 minute. **B.** The serum blocking of Y501-RBD to ACE2 with a similar dilution and procedure as A. **C.** The difference of binding of ACE2 between N501-RBD and Y501-RBD at different dilution of serum.

In November 2020, three therapeutic antibodies specific targeting RBD of SARS-COV-2, Bamlanivimab (LY-CoV555) from Eli Lilly, Casirivimab (REGN10933) and Imdevimab (REGN10987) from Regeneron were authorized for emergency use by the United States Food and Drug Administration (Bamlanivimab 2020; casirivimab and imdevimab 2020). It is important to find out whether Y501-RBD binding to ACE2 is still efficiently blocked or neutralized by these antibodies. Bamlanivimab was derived from a B cell from a convalescent patients and found to block efficiently the binding of N501-RBD to ACE2 (Chen et al. 2021). From the structural information of the online version, Bamlanivimab binds to a region on RBD close to Y501 (Jones et al. 2020). To find out whether Bamlanivimab would also block binding of Y501-RBD to ACE, we collected some residual of Bamlanivimab after infusion of COVID-19 patients and carried out binding assays of this antibody against either N501-RBD or Y501-RBD. To our delight, this antibody bound to N501-RBD or the variant Y501-RBD with a very similar binding affinity of 0.8737 nM (**Fig.4A**) and 0.8013 nM (**Fig.4B**) respectively. Thus, our data suggest that Bamlanivimab should be still effective for COVID-19 patients contracting either with wildtype SARS-COV-2 or with the 501Y.V1 variant, which first was found in the United Kingdom and spreads to the rest of the world recently. Of note, the binding affinity between the antibody and the United Kingdom variant Y501-RBD (0.8013 nM) is even lower than Y501-RBD and ACE2 (0.566 nM), this may indicate that much higher concentration of Bamlanivimab is needed to reach the similar therapeutic effect to treat COVID-19 patients with the United Kingdom variant compared to those with wildtype SARS-COV-2. Furthermore, for both variants from South Africa and Brazil, two additional mutations (K417N and E484K) happen within the receptor binding surface of RBD. Most strikingly, the side-chain of E484 of RBD and the side-chain of an Arg (R50) from Bamlanivimab (LY-CoV555) were found to form a salt bridge in the complex structure (Jones et al. 2020). The conversion of E484 to K484 (in the variants) could dramatically change the interaction mode between RBD and the antibody, leading, perhaps, to a dramatic reduction or loss of interaction, thus this monoclonal may be less useful in treatment of COVID-19 patients with either South Africa or Brazil variant.

**Figure 4.**
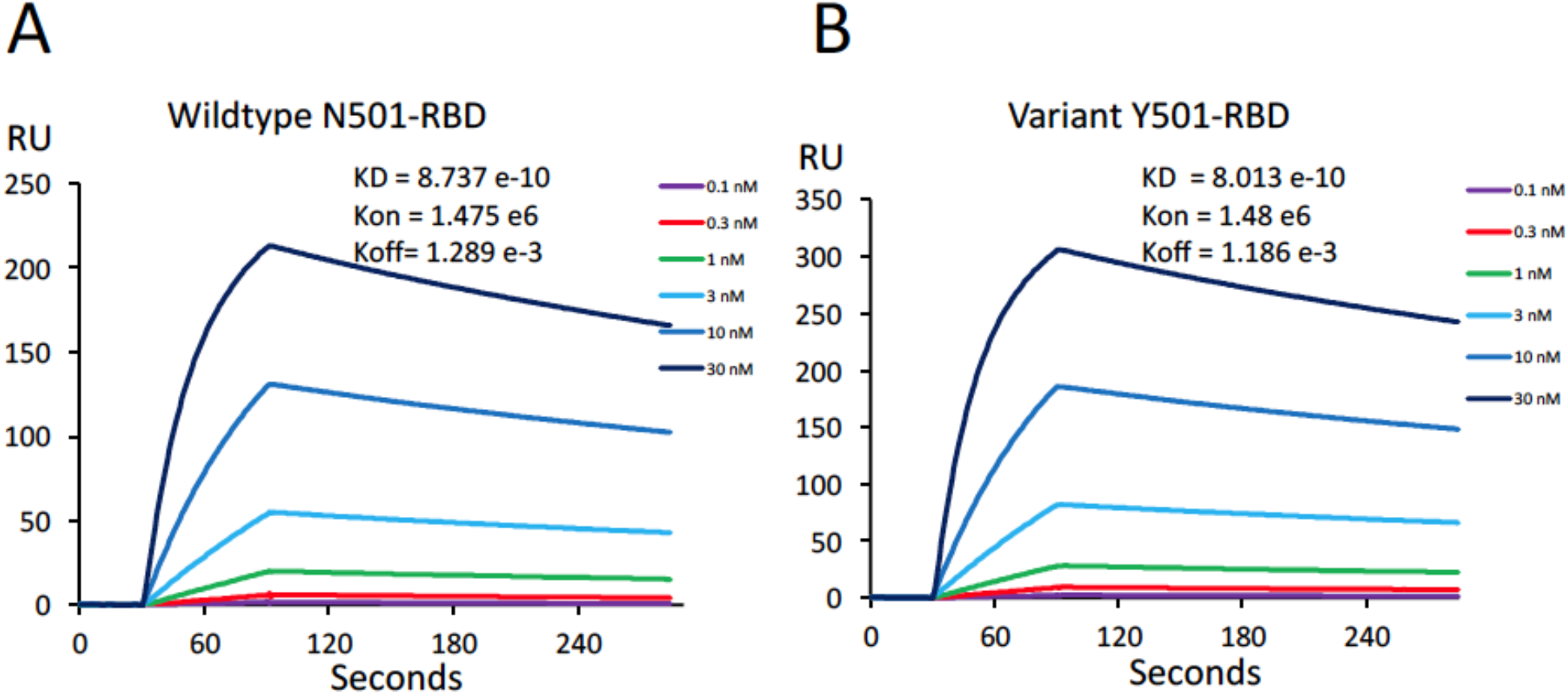
The binding affinities of N501-RBD and Y501-RBD to Bamlanivimab. **A.** The binding of N501-RBD to Bamlanivimab, immobilized Bamlanivimab with injection of different concentration of N501-RBD, **B.** The binding of Y501-RBD to Bamlanivimab, same as A.

The COVID-19 pandemic could be one of major disasters in human being history, and it is still out of control with limited means to contain it. Any mutation associated with an increased rate of infection could be a warning sign of potential escape of the virus from existing therapeutic options and pose challenges to both the available vaccines and therapeutic antibodies. Luckily, the mutation rate of the virus is low. Nevertheless, mutations in critical sites still could threat the efficacy of vaccines and therapeutic neutralizing antibodies. It should be predictable that the memory in both B cell and T cell generated by vaccines could be protective for the slowly evolving SARS-COV-2. Even the loss of efficacy of broadly neutralizing antibodies against the RBD regions with occasional escaping variants, the memory T cells with completely different protection mechanism through conserved linear antigens derived from SARS-COV-2 could build up a strong defense firewall against newly emerging and slowly evolved SARS-COV-2 variants. It is believed that broad vaccinations with two mRNA based vaccines from both Pfizer-BioNTech and Moderna or others will build up herd immunity to beat SARS-COV-2. Nevertheless, mutations within the surface of the ACE2 binding region on RBD, such as the United Kingdom variant, drastically increase the binding affinity (~10 times fold) between RBD and ACE2, it may require much higher concentration of the antibody to treat COVID-19 patients with the United Kingdom variant compared to patients with wildtype though it does not affect the binding of therapeutic antibody Bamlanivimab. On the other hand, some additional mutations within the RBD regions, such as K417N and E484K besides N501Y within both the South Africa and Brazil variants, could cause loss of efficacy of therapeutic antibodies, which are specific for this region. From this regard, therapeutic antibodies to treat COVID-19 patients should be adjusted for new emerging variants accordingly. In conclusion, from our current data, Pfizer-BioNTech vaccine and therapeutic antibody, Bamlanivimab from Eli Lilly, could protect people from wildtype SARS-COV-2 and the United Kingdom variant (501Y.V1).

## Methods

### Protein expression, purification, and binding assays

Human ACE2 ectodomain (1-615 aa) was cloned to pcDNA3 plasmid with 6-histidine tag and a BirA tag at the C-terminal. The protein was transiently expressed in 293F cells and purified by nickel column. SARS-CoV-2 spike RBD (319 aa-541 aa) was cloned to pcDNA3 plasmid. N501Y mutant was created by quick change mutagenesis and was verified by DNA sequencing. The wild type and mutant RBD protein was expressed in the same way as the ACE2. The protein was applied to a Superdex-200 Gel-filtration size column for further purification. BirA tagged ACE2 protein was biotinylated at the BirA tag using the BirA enzyme. The biotinylated ACE2 protein was collected at the right size on the sizing column.

### Affinity measurement between wildtype or mutant RBD and ACE2 using Biacore

Affinity measurement was carried out on the Biacore T200. Biotinylated ACE2 was immobilized on the surface of streptavidin chip on two separate channels for either wild type or N501Y mutant RBD measurement. A blank channel was used as background subtraction. Serial dilution of wild type RBD or N501Y RBD was injected to measure the affinity. The affinity was calculated by BIA evaluation software.

### ELISA measurement of human serum spike receptor binding domain antibody titer

RBD was coated on the ELISA plate in cold room overnight. The ELISA plate was then blocked with PBS with 5% FBS. Serial dilution of human serum was incubated on the plate at room temperature for 1 hour. The bound antibody was detected with AP conjugated goat-anti- human IgG antibody.

### Serum competing experiments

Same amount of wildtype RBD or N501Y mutant RBD was coated on two separate channels by the amine coupling method on the same CM5 chip. A third channel on the same chip was used as background control for signal subtraction. Serial dilution of human serum was injected with the same amount to two RBD channels (**Fig. S2**). When finishing injecting human serum, 20 μl of Biacore running buffer was injected to observe the serum antibody dissociation from RBD. 5 μg/ml recombinant human ACE2 was then injected to calculate the ACE2 binding to the unoccupied RBD. The BIAevaluation software was used to align the ACE2 binding to wildtype or mutant RBD coated channels for the series of serum dilutions. The ACE2 injection time point was aligned as baseline for the measurement of bound ACE2 to the wildtype (Fig 1E) or mutant RBD (Fig 1F) channel for each serum dilution. The difference of bound ACE2 in wildtype or mutant RBD channel was then used to plot Fig 1G.

### Eli Lily LY-CoV555 monoclonal antibody affinity measurement

The affinity was measured on Biacore T200. Eli Lily CoV555 mAb was coated on one channel of CM5 chip by amine coupling. A blank channel on the same chip was used as subtraction control. Serial dilution of wild type or N501Y mutant RBD from 0.1 nM to 30 nM was injected. The bound RBD was eluted with 10 mM glycine pH1.7 before injecting next cycle of RBD. The affinity was calculated by Biacore evaluation software.

### Protein docking

Protein docking studies were carried out using HADDock 2.2 server(Van Zundert et al. 2016). Crystal structure of SARS-CoV-2 Spike receptor-binding domain bound with ACE2 (PDB ID: 6M0J) was used for docking. To docking RBD N501Y mutant to ACE2, the RBD amino acid at position 501 was mutant from Asparagine (N) to Tyrosine (Y) by Coot software(Emsley and Cowtan 2004b). For all alterative conformation residues, one of the conformations in the structure was deleted manually to meet the docking server criteria. The ACE2 and RBD N501 structure were saved as a PDB files separately. The major contacts residues in the structure (PDB ID: 6M0J) (hydrogen bond and salt bridge) were selected as restrain residues for docking (Q24, D30, E35, E37, D38, Y41, Q42 and Y83 for ACE2 protein; K417, N487, Q493, Y449, and Y505 for RBD N501Y protein). The best model from the docking was selected for the next analysis. The structures were shown using PyMOL.

## Acknowledgements

We thank National Jewish Health for supports, People in Kappler/Marrack groups for help. H.L is partially supported by NIH grant (GM135421 to G.Z,) and NB Life Laboratory LLC. We thank Biobank Committee of National Jewish Health, Pearlanne Zelarney, Dr. Anthony Gerber, Joy Zimmer etc. for emerging approval, National Jewish Health Biobank for samples, Drs. Marco Ramirez-Gama and J. Tod Olin for contributions. We also thank the department of health, the state of Colorado, for authorization of usage of waste of therapeutic antibodies left after COVID-19 patient infusions.

## Contributions

H.L., P.M., and G.Z. for designing; H.L., Q.Z. for main experiments; P.W., Z.C., K.A., J.Y., W.D., S.P., G.D., L.R., S.F. for some experiments; C.J., B.L. J.K. for some data analysis; H.L., Q.Z., P.M., G.Z. for final data analysis and writing up.

## Supplementary material

**Figure S1.**
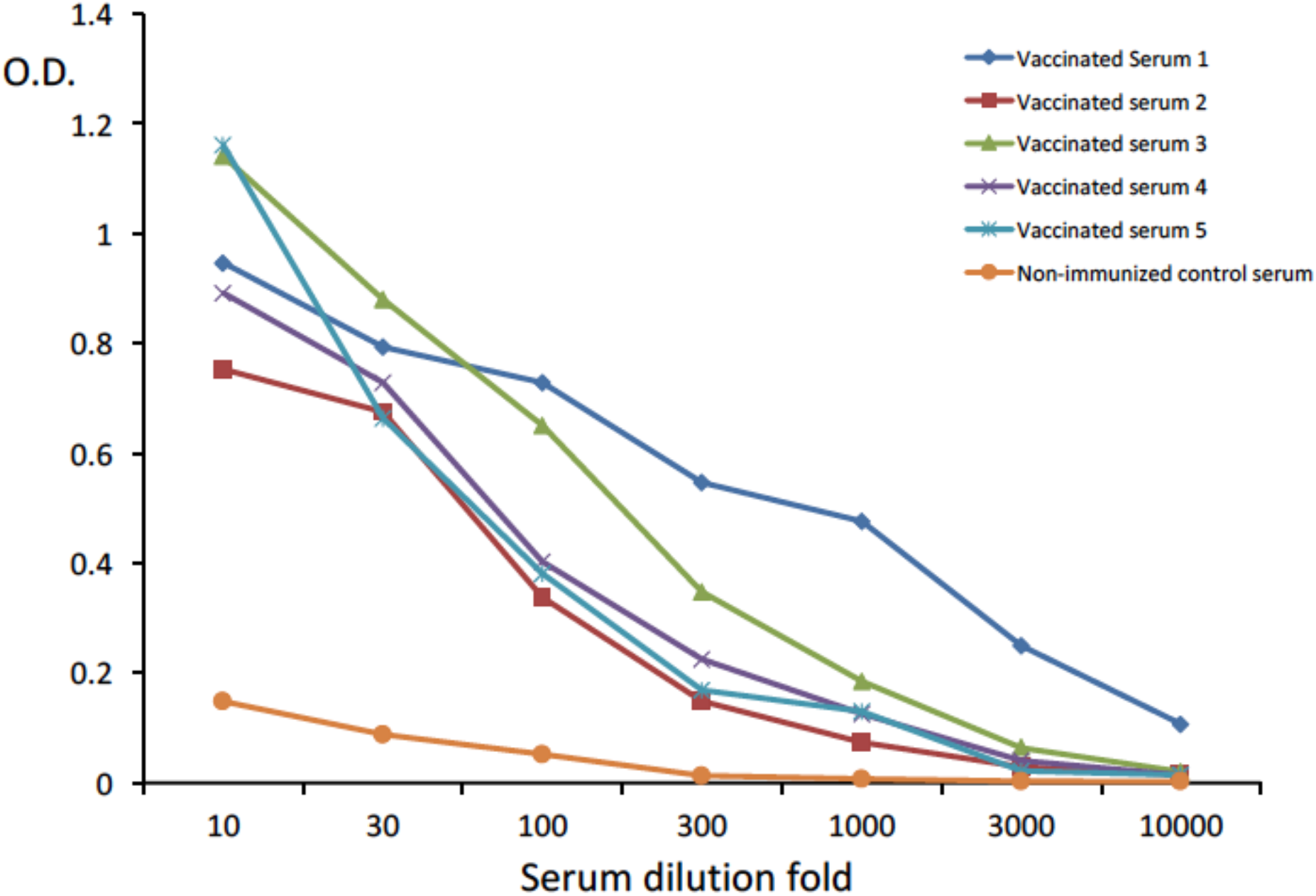
The titers of antibodies of sera from five individuals immunized with Pfizer-BioNTech vaccine. Blood samples are obtained from National Jewish Health Biobank.

**Figure S2.**
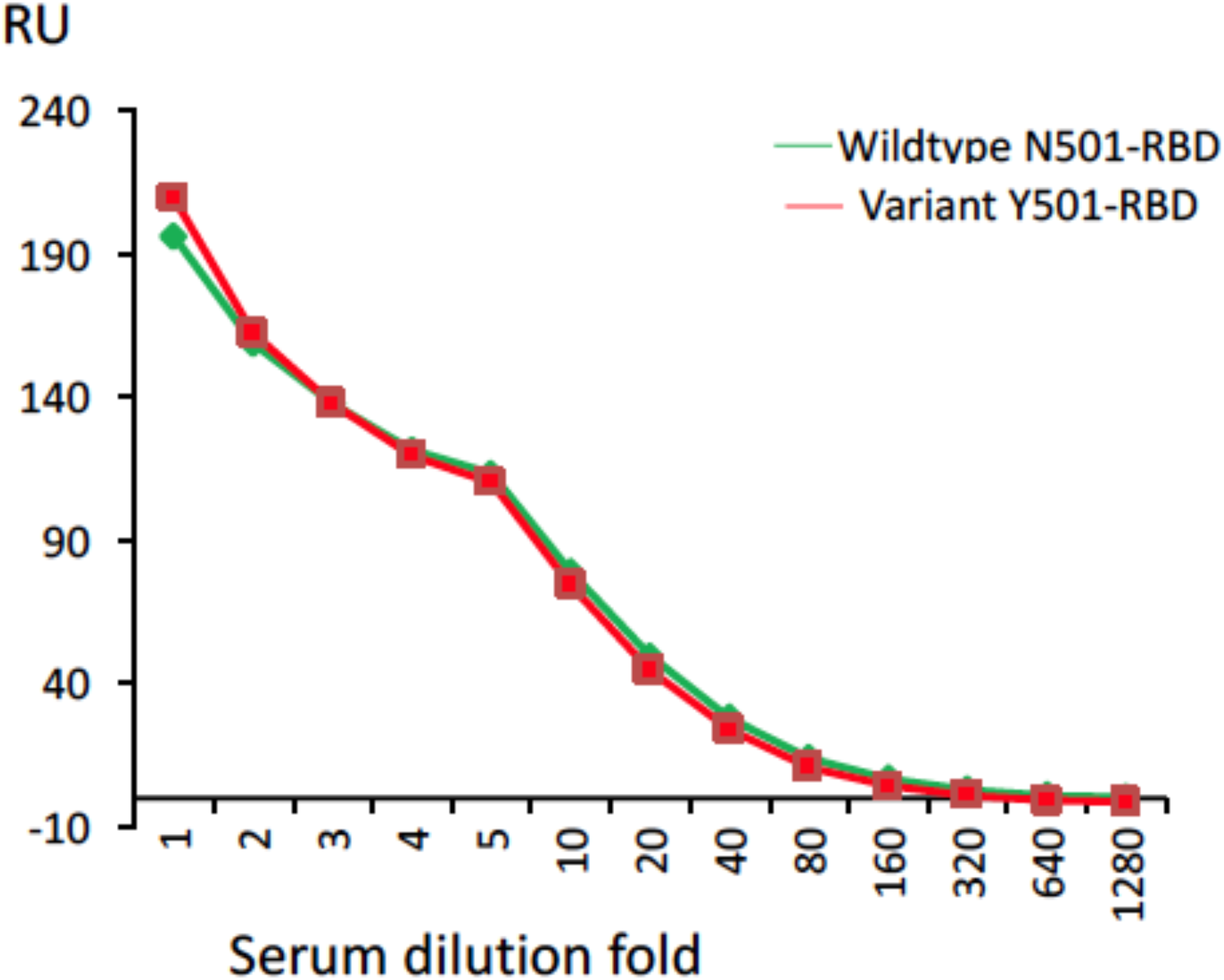
Similar amounts of antibodies bind to the two channels with N501-RBD and Y501-RBD respectively.

